# G-Quadruplex Mediated c-myc Specific Downregulation: A Unique Pathway of the Anticancer Action of Immunomodulator Drugs

**DOI:** 10.1101/2024.05.21.595106

**Authors:** Sunipa Sarkar, Akash Chatterjee, Subhojit Paul, Asim Bisoi, Prosenjit Sen, Prashant Chandra Singh

## Abstract

Hydroxychloroquine (HCQ), and chloroquine (CQ) are in the preclinical trial stage for cancer along with their active application in autoimmune diseases and malaria. One of the critical hallmarks of cancer cells is the elevated expression of various oncogenes which promote cancer progression and contribute to poor prognosis. The upstream of the promoter region of these oncogenes often exhibits a G-quadruplex (G4) DNA structure which regulates the gene expression. Hence, targeting G4 structure has emerged as a promising therapeutic strategy for cancer. In this study, the recognition of HCQ and CQ with the G4 structure of different oncogenes and its effect on gene regulation has been explored by a combination of various biophysical and *in-vitro* and *in-vivo* biological methods. This study depicts that HCQ and CQ downregulate the c-myc oncogene transcription significantly in a G4-dependent manner compared to other oncogenes. The different biophysical techniques and molecular dynamics simulation studies illustrate that these drug molecules stack predominately at the terminal of the c-myc G4 and the binding of these molecules stabilizes c-myc G4 significantly higher than the G4 structure of other oncogenes. The *in-vitro* cell data exhibit a notable reduction in both c-myc mRNA and protein levels in a triple-negative breast cancer cell line following HCQ treatment. The pre-clinical breast cancer mouse model *in-vivo* data also indicate that HCQ reduces tumor growth through the downregulation of the c-myc oncogene. Simultaneously, HCQ also enhances the therapeutic efficacy of standard chemotherapeutic agents to be a potential candidate for combination therapy. This work demonstrates the alternative strategy of anticancer action of widely used drugs by specifically downregulating the c-myc oncogene in a G4-dependent manner.

## Introduction

The prevalence of cancer constitutes a formidable threat to human health, with chemotherapy comprising a frontline therapeutic intervention.^1^ However, extant chemotherapeutic regimens frequently engender significant side effects and can elicit acquired resistance, precipitating disease relapse.^1^ The oncogenesis is intricately linked to genetic aberrations in critical regulatory genes, notably the aberrant expression of various oncogenes that propagate dysregulated signals leading to cell proliferation, migration, increased invasion into the surrounding matrix, etc.^2, 3^ Elucidation of novel chemotherapeutic targets and strategies to mitigate treatment resistance remains imperative to improve clinical outcomes for cancer patients. The elucidation of innovative chemotherapeutic targets and the development of approaches to overcome treatment resistance are of critical importance in order to enhance therapeutic efficacy and clinical outcomes among cancer patients.^1, 2^ c-myc emerges as a pivotal oncogene, exhibiting upregulation in approximately 70% of cancer instances.^4, 5^ Directly addressing the overexpressed c-myc protein poses a formidable challenge owing to the absence of viable small molecule binding sites and its limited lifespan.^6, 7^ Consequently, efforts have been redirected towards the exploration of alternative avenues in cancer treatment, focusing on targeting specific DNA secondary structures associated with oncogenes, with particular emphasis on the extensively overexpressed c-myc.^8–11^

G-quadruplexes (G4) are secondary DNA structures formed by the folding of guanine-rich DNA sequences in the presence of monovalent cations.^12, 13^ G4 structures are stabilized through the Hoogsteen-type hydrogen bond and π stacking between the guanine nucleobases. Several genomic sequences having guanine-rich domains are found to be more thermodynamically stable than the double-strand DNA and these sequences can directly form G4 structure.^14^ The formation of G4 structures has been identified in the promoter region of several oncogenes through computational and experimental studies.^15–19^ Stabilizing these G4 structures within oncogene promoters has been demonstrated to reduce transcriptional rates, offering a novel avenue for cancer therapy.^20–24^ Despite concerted efforts to develop small molecule ligands capable of stabilizing G-quadruplex structures within oncogene promoter regions, no compounds have yet garnered clinical approval as targeted antineoplastic agents, ostensibly due to lingering uncertainties regarding their compatibility with biological systems and suboptimal pharmacological properties.^25–30^ Therefore, it is imperative to investigate the gene regulatory effects of the drug candidates currently undergoing clinical trials, specifically their interactions with G4 structures. Such investigations are crucial as these molecules hold potential as viable candidates for future cancer therapies.

Hydroxychloroquine (HCQ, Figure 1a) and chloroquine (CQ, Figure 1a) are important immunomodulator drugs that have been extensively used for the treatment of rheumatoid, systematic lupus erythematosus, and malaria.^31–33^ Recently, it has been reported that HCQ and CQ may affect the cancer cells and may increase the tumor sensitivity to existing cancer treatments.^34–36^ Moreover, numerous clinical trials have been initiated to evaluate the therapeutic potential of both HCQ and CQ as novel targeted agents against a diverse array of cancer histologies, with over thirty studies currently registered across all phases of clinical evaluation.^35, 37^ There are few studies in which the plausible action of HCQ and CQ on different cancer cell lines have been discussed.^38–42^ Inhibition of autophagy is one of the most studied anticancer actions of HCQ and CQ.^38^ It has been proposed that these drugs get protonated upon entering lysozyme which gets them trapped in the acidic lysosome and inhibits the lysosomal degradative enzymes.^38^ However, recent studies suggest that CQ seems to transiently perturb lysosome integrity and function rather than suppressing an integrated autophagy process as proposed earlier.^39^ The other few proposed actions of the CQ are the inhibition of kappa B and CXCL2/CXCl4 signaling pathways.^40–42^ Apart from these studies, the role of HCQ in tumor immunotherapy has been investigated, however, it has been proposed that HCQ decreases the benefit of anti-PD1-immune checkpoint blockade in tumor immunotherapy.^43, 44^ Hence, it can be speculated that the antineoplastic efficacy of immunomodulator drugs HCQ and CQ may have another selective biological target. While a limited number of studies have explored the putative antineoplastic efficacy of HCQ and CQ across various cancer cell lines, the precise molecular mechanisms underpinning their activity remain incompletely defined.^38–44^

**Figure 1:**
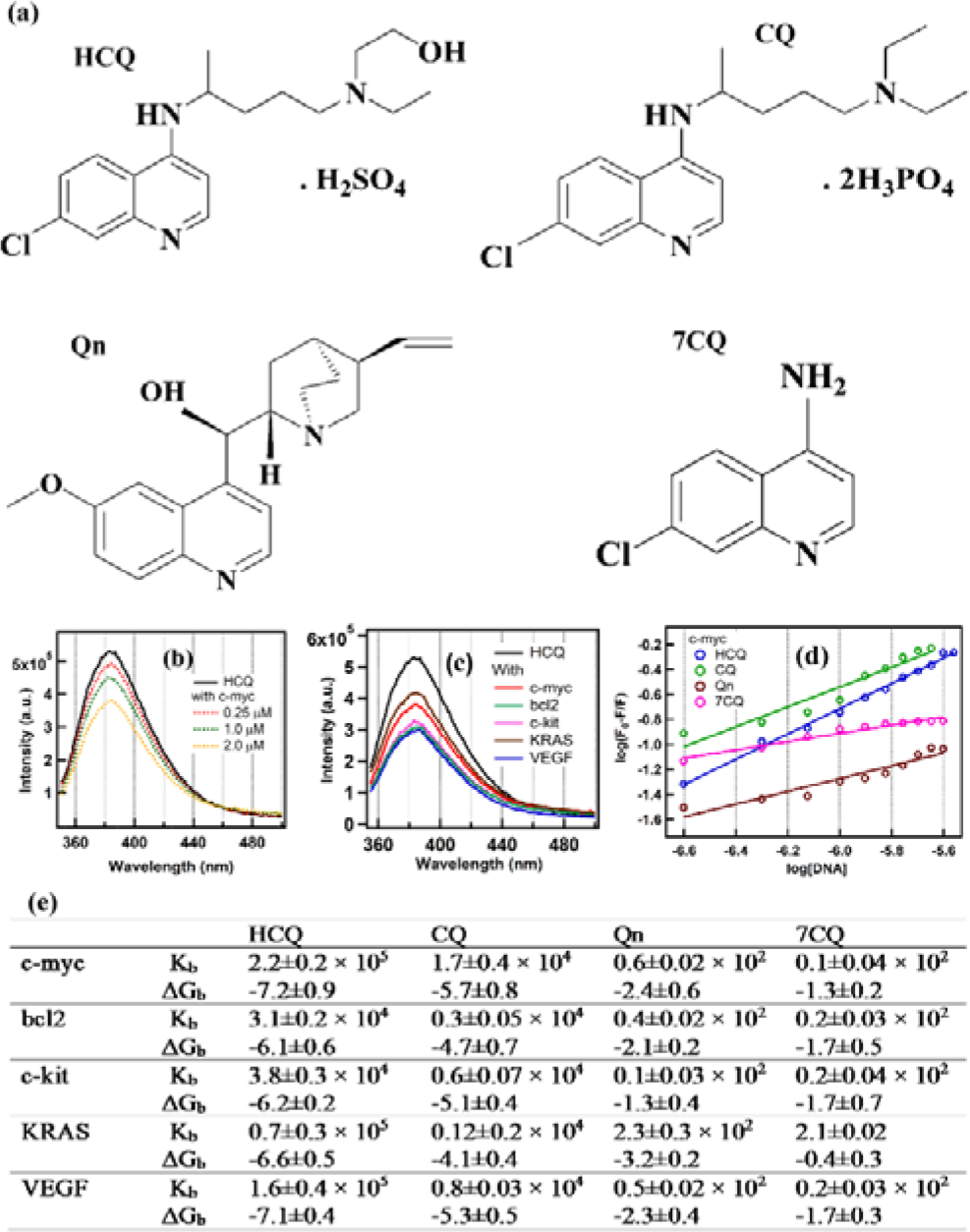
(a) The chemical structure of drug molecules HCQ, CQ, and their structural analogs Qn and 7CQ, respectively. (b) The emission spectra of HCQ in buffer and different concentrations of c-myc DNA. (c) The emission spectra of HCQ in buffer and different G4 DNA sequences. (d) The plot of the change in the intensity of the emission maxima of HCQ, CQ, and its analogs Qn and 7CQ with respect to the concentration of c-myc DNA. The fitting of the plot has been performed using equation 1 to get the binding constant (K_b_) of the drug with DNA sequences (e) The values of K_b_ and binding free energy (ΔG_b,_ kcal/mole) of drugs with gene sequences at 25L.

Recently, the possibility of the binding of HCQ with G4 has been illustrated by our group using the spectroscopic and molecular simulation methods.^45^ However, despite the evident therapeutic imperative to selectively target aberrantly overexpressed oncogenes, the antineoplastic efficacy of immunomodulator agents against such targets remains largely unexplored. Elucidating the precise molecular mechanisms by which this class of pharmacologic impacts overactivated oncogenic signaling cascades would augment their clinical development as targeted chemotherapeutics. In this study, we have established that immunomodulator HCQ and CQ significantly reduced the transcriptional activity of the c-myc gene predominantly stabilizing the G4 in upstream of the promoter region of the gene and significantly restricting the progression of cancer. Firstly, the spectroscopic analysis demonstrated preferential stabilization of the G4 structure within the c-myc promoter sequence by HCQ and CQ compared to those formed in other oncogenic promoter regions. Subsequent investigation using NMR spectroscopy and molecular dynamics simulations enabled further characterization of the precise binding modality underpinning HCQ-c-myc G4 interactions. Different *in vitro* biochemical experiments demonstrate significant downregulation of the c-myc gene in a G4-dependent manner by HCQ than other genes. The *in-vitro* findings of specific downregulation of the c-myc gene by HCQ have been also corroborated by employing a pre-clinical breast cancer murine model. Finally, our analysis additionally revealed the HCQ potentiation of c-myc transcriptional suppressive activity when administered as adjunct therapy with the classical chemotherapeutic agent, doxorubicin, suggesting potent synergy between these agents. These findings elucidate a novel therapeutic paradigm for targeting triple negative breast cancer via immunomodulatory agents such as HCQ and CQ, predicated upon selective transcriptional repression of the c-myc proto-oncogene through stabilization of G4 structures. This approach represents a promising alternative strategy to improve clinical outcomes in this aggressive disease subtype.

## Materials and methods

### Chemicals

High-pressure liquid chromatography grade pure DNA sequences of c-myc, bcl2, c-kit, KRAS, VEGF, and duplex DNA were purchased from Sigma Aldrich. HCQ, CQ, quinine (Qn, Figure1a), 4-amino-7-chloroquine (7CQ, Figure1a), and KCl were procured from Sigma Aldrich. All the DNA samples were dissolved in 10 mM Tris-HCl buffer (pH=7.4) and annealed in the 2 mM KCl salt by increasing the temperature up to 90_ followed by cooling at room temperature. The annealed samples were kept in a refrigerator at 4_ for further use.

### Fluorescence measurements

The emission spectra of the immunomodulator drugs dissolved in Tris-HCl buffer (pH=7.4) are measured in the absence and presence of DNA samples using Fluoromax-4 (Horiba Scientific) instruments at 25_. The slit width of the measurement is fixed at 2 nm. The binding constant of drug molecules with different sequences of DNA has been estimated by measuring the fluorescence of these drugs with increasing concentrations of different DNA sequences. The binding constant was calculated using the modified Stern-Volmer equation shown in equation (1).

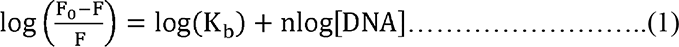

Here, F_0_ and F correspond to the fluorescence intensity of the drug in the absence and presence of DNA. K_b_ and n represent the binding constant and stoichiometry of the binding of drugs with different sequences of DNA. The binding free energy ΔG_b_ (kcal/mole) of drugs with DNA sequences was calculated at 25_ using equation 2.

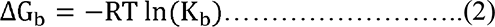

### Circular dichroism and melting temperature measurements

The CD spectra of the G4 structure of each sequence of DNA in the presence of drugs in the range of 220-310 nm were measured using JASCO (J1500) spectrometer to understand the effect of the drugs on the G4 structure of the different sequences of DNA. The drug-induced stabilization in the G4 structure of the different sequences has been estimated by measuring the melting temperature of the G4 structure of DNA in the absence and presence of drug molecules. The melting temperature of each G4 structure was estimated by measuring the temperature-dependent change in the ellipticity of the characteristic peak of the G4 structure (∼260 nm). The temperature of the system was changed from 10_ to 90_ in the interval of 3_.

### NMR measurement

The ^1^H-NMR measurements in the imino region of c-myc (100 µM) annealed in 2 mM KCl were measured in the presence of HCQ and 7CQ in H_2_O/D_2_O 90%/10% mixture to understand the binding modes of HCQ with c-myc. The NMR measurements were performed using a 600 MHz spectrometer at 25_. Each spectrum is an average of 10000 scans.

### Molecular dynamics (MD) simulation

MD simulation was conducted utilizing the parallel G4 structure of c-myc (1AXV.pdb)^46^, as described by the Amber99 force field with OL15 correction.^47^ Three distinct systems were prepared: c-myc without a drug, c-myc with HCQ, and c-myc with 7CQ. The parameters for the protonated form of HCQ^48^ and the neutral structure of 7CQ were adopted from the Generalized Amber Force Field (GAFF) using the Antechamber package.^49^ The solvation of the systems was achieved using TIP3P water molecules.^50^ For the first system, the G4 structure was solvated in a cubic box with sides measuring 6 nm, maintaining a 0.1 M salt concentration without a drug molecule. Subsequently, for the second system, six HCQ cations and six SO_4_^2-^ anions were introduced. For the third system, six 7CQ molecules were randomly inserted within the box. Ions (K^+^, Cl^-^) were employed to maintain a 0.1 M bulk concentration of salt and neutralize the negative charge of G4. The simulations were conducted using GROMACS-2016.3 software.^51^ Initially, all systems underwent energy minimization for 5000 steps using the steepest descent method, followed by position-restrained equilibration for 200 ps. Subsequently, NVT (300 K) and NPT (1 atm) simulations were performed for 1 ns each to properly equilibrate the solvent and ions. The production MD run spanned 1 µs for each system, employing a time step of 2 fs. The LINCS algorithm was utilized to constrain bond lengths between heavy atoms and hydrogen.^52^ Temperature and pressure were regulated through the velocity rescaling (V-rescale) thermostat^53^ (τ=0.1 ps) and Parrinello-Rahman pressure coupling^54^ (τ=2 ps), respectively. The cutoff radius for neighbor searching and nonbonded interactions was set at 1 nm, with the nonbonded pair list updated every 10 steps. The particle Mesh-Ewald method with a grid spacing of 0.12 nm and fourth-order interpolation was applied to account for long-range electrostatic interactions.^55^ Trajectory analysis utilized GROMACS tools, and snapshots were generated using Visual Molecular Dynamics software (VMD 1.9.3).^56^ The radial distribution function of the HCQ and 7CQ with different parts of G4 was calculated to get an idea of the interaction of these molecules with G4. The following atoms of the G4 have been considered to calculate the RDF of the quartet, loop, and the phosphate backbone of the G4 with drug molecules: (1) Electronegative atoms of the guanine bases for the quartet region, (2) the electronegative atoms of the T and A bases for the loop regions, (3) Phosphorus of the phosphate backbone have been selected for RDF calculations. The nitrogen of the quinoline ring of HCQ or 7CQ (denoted as N2) has been considered for the RDF calculation of the ring part of drug molecules whereas the side chain has been represented by the O and the N atoms of the sidechain for HCQ and 7CQ, respectively. The total interaction energy of the drug with quartet, loop, and phosphate backbone of G4 and its coulombic (Coul) and Lennard-Jones (LJ) terms has been calculated to get the idea of the energetics of binding of these drug molecules with G4. For the energy calculation, quartet: heavy atoms of the G bases of the rings without phosphate backbone, loop: heavy atoms of the A, and T bases without phosphate backbone, and phosphate: heavy atoms in the phosphate regions were selected.

### Cell culture

The human cancer cell lines MDA-MB-231 (human triple-negative breast cancer cell line), HeLa (human cervical cancer cell line), and murine breast cancer cell line 4T1 were acquired from the American Type Culture Collection and cultured in DMEM (Gibco) medium supplemented with 10% FBS and 100 units/ml penicillin-streptomycin (Invitrogen).

### Western blotting

To find out the expression of c-myc and bcl2 in cancer cells, 5 × 10^5^ cells were seeded per well in a 6-well plate. Cells were kept overnight in complete growth media to adhere. They were then treated either with different concentrations of HCQ (100 µM, 50 µM, 25 µM, and 12.5 µM) or with 50 µM CQ, Qn, 7CQ, and HCQ (dissolved in pH 7 buffer) for 48 hours. Cells were then lysed using Laemmli buffer. The lysate was heated at 95 °C for 5 min. SDS-PAGE was performed to separate the proteins and transfer them onto the PVDF membrane, following blocking in 5% BSA in TBS buffer. The membrane was incubated overnight at 4 °C with primary antibodies for specific proteins in 3% BSA in TBS. A secondary antibody was added and incubated for an hour after washing with TBS-T (TBS, 1% Tween20) three times. Then blots were developed using ECL. Densitometric analysis was done using ImageJ, and graphs were made using GraphPad Prism 8.

### Flow cytometry

To determine the expression of c-myc, cells treated with 50 µM of HCQ, CQ, Qn, 7CQ, (dissolved in pH 7 buffer) for 48 hrs were washed with PBS. Then the cells were permeabilized with 1% Triton X-100. Next, cells were incubated with c-myc primary antibody for 1 hr, followed by 45 min incubation with Alexa-488 conjugated corresponding secondary antibody. Rabbit IgG was used as isotype control. Cells were then subjected to flow cytometry analysis (BD FACS AriaIII). Acquired data were analyzed using FlowJo (version 8).

### Quantitative real-time PCR (qRT-PCR)

To evaluate the influence of HCQ on the mRNA expression of c-myc, bcl2, c-kit, KRAS, VEGF, and HK-2 in cancer cells (Table S1), cells were seeded at a density of 5 × 10^6^ cells per 35 mm dish. Next cells were treated with different doses (100 µM, 50 µM, 25 µM, and 12.5 µM) of HCQ or fixed 50µM of HCQ as per desired experiments. Subsequently, total cellular RNA was extracted using the TRIZOL reagent (Life Technologies). cDNA was then prepared from extracted total RNA using the Super-Script III First-Strand cDNA Synthesis System from Invitrogen, following the manufacturer’s guidelines. The resulting cDNA was utilized for quantitative real-time PCR, employing the SYBR Green Master Mix (Applied Biosystems), performed on the Step One plus Real-Time PCR System. Data normalization was accomplished using GAPDH as the reference gene, and each experimental sample underwent triplicate analyses for precision.

Relative expression levels were computed using the 2^-ΔΔCt^ method. Primer sequences are provided in the supplementary table.

### Immunofluorescence

Cells were seeded onto hydrofluoric acid-etched coverslips (35%). Subsequently, cells received treatment with either 50µM or 100µM of HCQ. Fixation was carried out using 4% paraformaldehyde for 30 minutes, followed by a 10-minute permeabilization step using 0.1% Triton X-100. After three washes with PBS, cells were blocked with 5% BSA for 1 hour. Overnight incubation with c-myc antibody followed, and cells were then subjected to a 1-hour treatment with Alexa Fluor 488-conjugated rabbit secondary antibody post-washing. Slides were exposed to DAPI for 10-20 minutes, followed by additional PBS washing. Finally, cell imaging was performed using a Carl Zeiss laser scanning confocal microscope.

### Cell viability assay

To assess cell viability, MDA-MB-231 cells (Human triple-negative breast cancer) were seeded at a density of 8 × 10^3^ cells per well in 96-well plates. After overnight incubation in complete growth media (DMEM CM), the cells were treated with one of the following: 50 µM of HCQ, 1.5 µM of DOX, or a combination of both compounds. These treatments were administered for 72 hours after which MTT reagent was added. Following a suitable incubation period, the formazan crystals that developed were dissolved in DMSO, and the absorbance was measured at 570 nm to determine cell viability. The resulting data were analyzed, and cell viability graphs were generated using GraphPad Prism 8.

### Migration assay

MDA-MB-231 cells were seeded onto 35-mm dishes, permitting them to attain 80–85% confluence. Subsequently, the cells underwent treatment with HCQ (50µM), DOX (1.5µM), or a combination of both. On the same day, uniform scratch lines were created using a micro tip. After a 48-hour interval, images were captured to assess migratory changes. The migratory capacity of cells was quantified by contrasting them with the baseline 0-hour treatment.

### Invasion assay

Cells were equally seeded onto Matrigel-coated Transwell chambers. After treatment with HCQ (50µM), DOX (1.5µM), or a combination, the upper chamber contained a serum-free medium while FBS-containing DMEM was in the lower chamber. Following 48 hours of incubation, cells that invaded the lower membrane surface were stained with crystal violet solution. After removing upper surface cells, the total invaded cells were counted in 10 randomly chosen fields using a bright-field microscope.

### Animal studies

Balb/c mice were purchased from W.B. Livestock Dev. Corpn. Ltd. All experiments were performed following the institutional guidelines. 6-8-week old BALB/c mice were subcutaneously injected (1x10^6^ 4T1 cells re-suspended in PBS buffer) in the lateral flank. Mice (n=5 per group) were then divided into groups and were then subjected to control (saline), oral gavage of either 10mg/kgbdwt or 50mg/kgbdwt HCQ, 5mg/kg of DOX treatment via tail vein injection, or a combination of both HCQ and DOX treatment were carried on the one-day interval from the 5^th^ day until the day of sacrifice. Tumor size was measured post-sacrifice at day 25 post-tumor cell injection. Tumors were removed weighed and compared between the groups. The tumor tissue was homogenized in a lysis buffer to extract proteins. Western blot analysis was then performed to measure the c-myc levels in different tumor groups.

## Results

### Binding of HCQ and its analogs with G4 DNA sequences of different genes

Figure 1b illustrates the emission spectra of HCQ in buffer, with varying concentrations of G4 DNA sequence of the promoter region of c-myc. The observed successive decrease in the fluorescence intensity of HCQ following the addition of c-myc G4 DNA suggests a plausible interaction between HCQ and the G4 DNA sequence of the c-myc gene, supporting the notion that the binding of HCQ with DNA results in the quenching of its fluorescence.^45^ The fluorescence of HCQ exhibited a decrement upon the addition of other G4-forming DNA sequences (bcl2, c-kit, KRAS, and VEGF) concurrently with c-myc (Figure 1c). This observation suggests that HCQ can interact with all these G4-forming DNA sequences. The fluorescence data for CQ, Qn, and 7CQ similarly demonstrate a reduction in fluorescence intensity upon the addition of these G4-forming sequences (Figure S1). Notably, HCQ and CQ manifest more pronounced reductions in fluorescence intensity upon the addition of G4 DNA sequences compared to their respective analogs, Qn and 7CQ. This observation implies that the binding propensities of the drugs HCQ and CQ with the G4 DNA sequences may differ from their analogs Qn and 7CQ. Therefore, the determination of binding constants (K_b_) and binding free energy (ΔG_b_) for all the molecules interacting with different G4-forming DNA sequences was executed through the assessment of fluorescence changes induced by the addition of DNA sequences and utilizing the modified Stern-Volmer method (equation 1), as depicted in Figures 1 d, e and S2. The calculated K_b_ and ΔG_b_ values indicate that the binding affinity of HCQ and CQ for any G4-forming DNA sequence is relatively higher compared to Qn and 7CQ. The structural configuration of HCQ and CQ closely resembles that of 7CQ, differing primarily in the presence of charge on the ring and the side chain. In contrast, Qn shares the quinoline ring structure but features distinct substitutions at the 4^th^ and 7^th^ positions compared to other synthetic analogs. Structural and binding correlation analysis suggests that the charge and side chain of HCQ and CQ are crucial for facilitating their interaction with G4 DNA sequences.

### Drug-induced structural stability of G4 associated with different genes

To assess the impact of drug molecules on the structural aspects of G4, CD measurements were performed for all G4 DNA sequences in both the absence and presence of drugs (Figure 2a). Consistent with prior studies, CD spectra of G4 DNA structures formed in 2 mM KCl displayed positive ellipticity peaks at 265 nm and negative ellipticity at 245 nm, indicative of the parallel conformation of the G4.^57, 58^ The introduction of drug molecules resulted in marginal changes in the CD peak ellipticity without the emergence of new peaks, suggesting that drug binding does not induce significant structural alteration in the parallel G4 conformation across all sequences. For a quantitative assessment of ligand-induced G4 stabilization, the melting temperature of G4 was determined in the presence and absence of drug molecules by monitoring temperature-dependent changes in the characteristic 260 nm CD peak of the G4 (Figures 2b, c, and S3). The data revealed a notable increase in the stabilization of G4-forming DNA sequences, particularly with HCQ and CQ (11-17°C), in contrast to Qn and 7CQ (0-5°C). This trend aligns with the binding data and emphasizes the crucial role of both the charge and side chain of HCQ and CQ in G4 stabilization. The presence of HCQ and CQ has minimal impact on the melting temperature of duplex DNA, indicating the selective nature of G4 stabilization by these drug molecules (Figure 2d). Noticeably, the HCQ/CQ-induced stability of G4 structures exhibited sequence dependence, with the highest change in melting temperature (ΔT_m_) for c-myc (∼17°C) compared to other G4 forming sequences such as bcl2 (7°C), c-kit (11°C), KRAS (12°C), and VEGF (9°C). The melting temperature data reveal the selective stabilization of HCQ and CQ to G4 structures located in the promoter regions of various genes, with the highest stabilization for the c-myc G4 relative to those in the promoters of other examined gene sequences.

**Figure 2:**
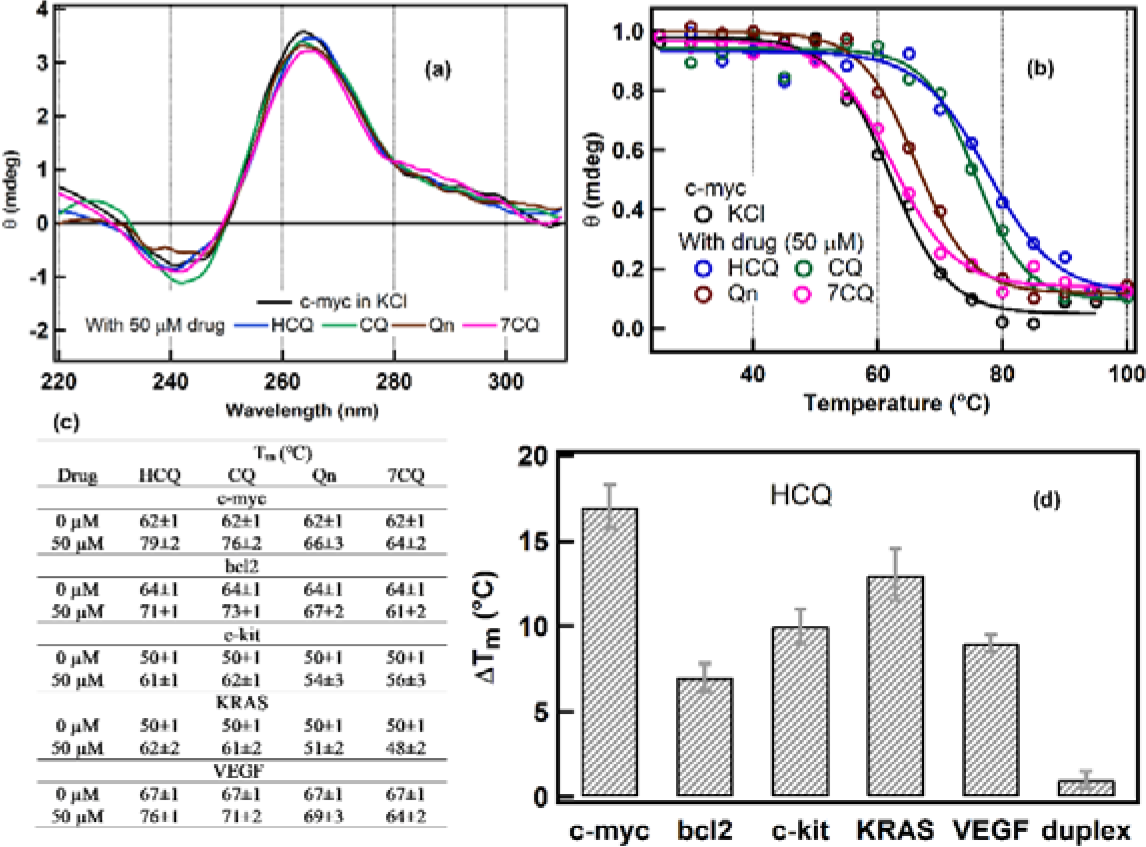
(a) The CD spectra of c-myc in KCl salt and in the presence of HCQ, CQ, Qn, and 7CQ drugs. (b) The melting curve of the c-myc in KCl and the presence of different drug molecules (c) The melting temperature of the G4 formed from the different sequences and the effect of the drug molecules on the melting temperature of these sequences. (d) The change in melting temperature (ΔT_m_=T_m_ of G4 in drug-T_m_ of G4 in buffer) of different G4 sequences in the presence of HCQ and its comparison with the duplex DNA sequences.

### Characterizing the binding of HCQ with c-myc G4 using ^1^H-NMR spectroscopy and MD simulation

The above-discussed data depict that the binding of HCQ and CQ provides significant stability to the c-myc G4 compared to G4 structures in other genes. Therefore, ^1^H-NMR experiments in the imino region (10-12 ppm) of the c-myc G4 have been performed to elucidate the binding of HCQ with c-myc G4 (Figure 3a). Consistent with earlier reports, the NMR spectra of c-myc G4 exhibited twelve well-resolved peaks, corresponding to the formation of three tetrads within the c-myc G4 (Figure 3b).^57–59^ Upon the addition of HCQ, the fast chemical exchange of protons in the c-myc G4 was perturbed, resulting in the appearance of new sets of peaks, indicative of the formation of the HCQ-c-myc G4 complex. The most significant perturbations in the chemical shift (CSP) of peaks were observed for G7, G11, and G16, which constitute the 5_ end of the c-myc G4, while interactions were less prominent with the 3_ end and the central tetrad of G4 (Figure 3c). Additionally, we conducted NMR experiments on the c-myc G4 with the addition of 7CQ (Figure S4). The change in NMR peaks of the G terads in the presence of 7CQ was similar to that observed with HCQ. A comparative analysis of the change in chemical shift data for c-myc in the case of HCQ and 7CQ suggests that the chromophoric quinoline group of HCQ is stacked predominately with the 5_ end of the c-myc G4.

**Figure 3:**
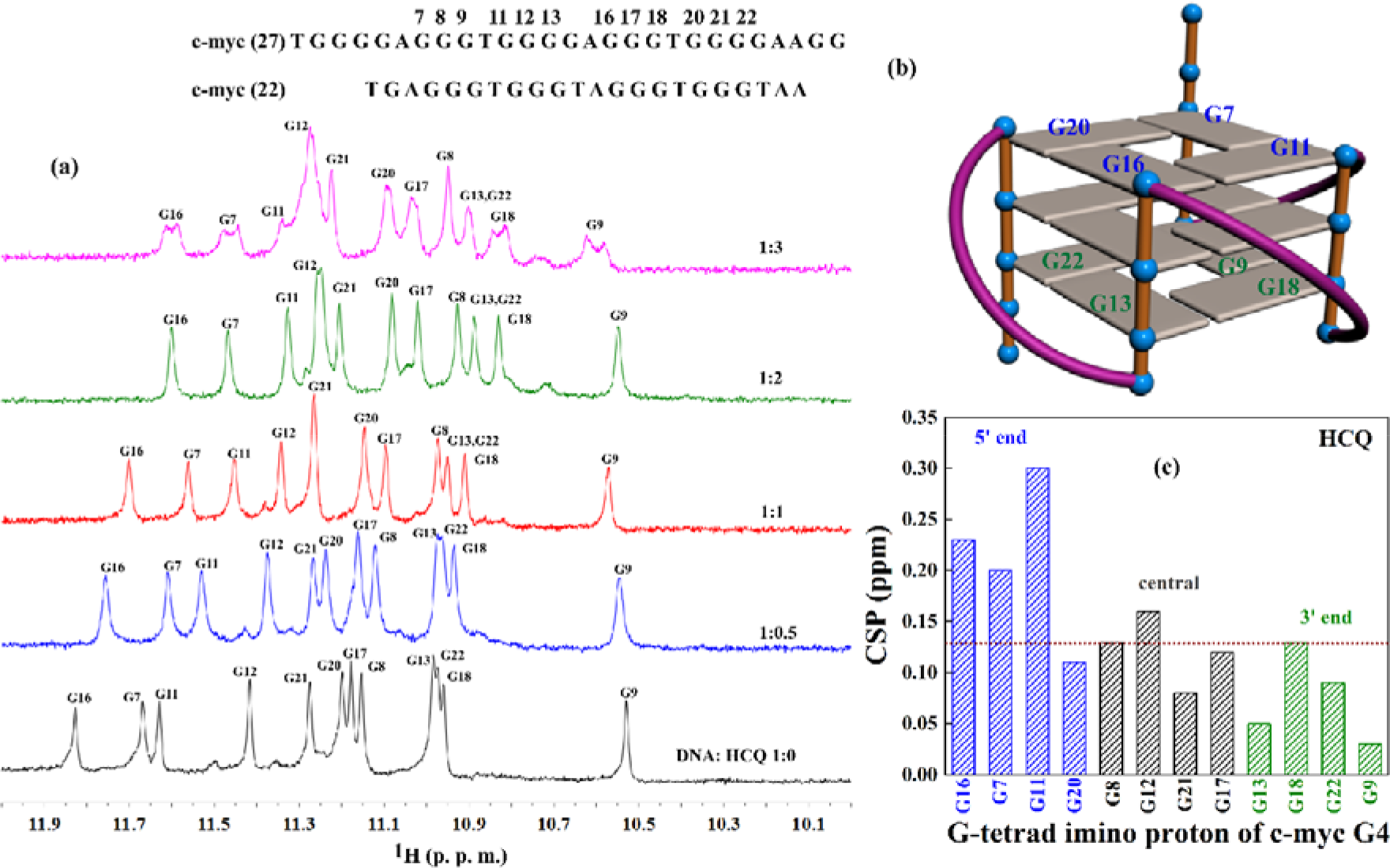
(a)^1^H NMR spectra representing the imino region of the G4 during the titration of c-myc with HCQ. The G4-imino protons of tetrad and the ratio of c-myc and HCQ are labeled in each spectrum. (b) The representation of the parallel structure of c-myc. The G4- tetrad at 5’ and 3’ terminals are marked in blue and green colors. (c) The plot of chemical shift perturbation for (CSP) each imino protons of c-myc G4-tetrads at 5’end (blue), central (black), and 3’ end (green) terminals. The CSP has been calculated by the subtraction of the position of each imino proton in the case of a 1:2 c-myc-HCQ complex with respect to the only c-myc case. The brown dashed line in Figure c indicates the one standard deviation above the average value of the CSP of all the cases.

Next, the 1 µs MD simulation of the G4 structure of c-myc in water and in the presence of HCQ and 7CQ has been performed to illustrate the binding of these drugs with G4. The time evolution of the root mean square deviation (RMSD) values of the G4 structure in the absence and presence of drugs is almost constant with slight fluctuation (Figure S5) indicating that the structure of G4 is intact on the binding of G4, in accordance with the CD data. The drug molecules, HCQ and 7CQ, possess two binding sites: one is defined as an aromatic ring having a quinoline ring (drug name_ring_), and the remaining part is considered as side chain (drug name_sidechain_).To delve deeper into the binding modes, the radial distribution function (RDF) of these two binding sites of HCQ and 7CQ with quartet, loop, and phosphate backbone regions of G4 has been calculated (Figures 4a-e). The RDF and the snapshot data indicate that both HCQ and 7CQ prefer to stack with the quartet and loop regions of G4 (Figure 4 a,b,d, and e), in agreement with the NMR results. However, the preference for the accommodation of the HCQ and 7CQ in the loop and quartet region is different as HCQ prefers to stack more with the loop than the quartet of G4 whereas 7CQ interacts with both, the loop and quartet of G4. The RDF of the HCQ and 7CQ with the phosphate backbone depict a distinct difference as the charged side chain of HCQ interacts strongly with the phosphate backbone than 7CQ (Figure 4c). The RDF and snapshot of the simulation indicate that the strong binding of the charged side group of the HCQ with the phosphate backbone cooperates with the stacking of its quinoline group with the quartets and loops of G4 which probably causes the strong binding of HCQ than 7CQ with G4.

**Figure 4.**
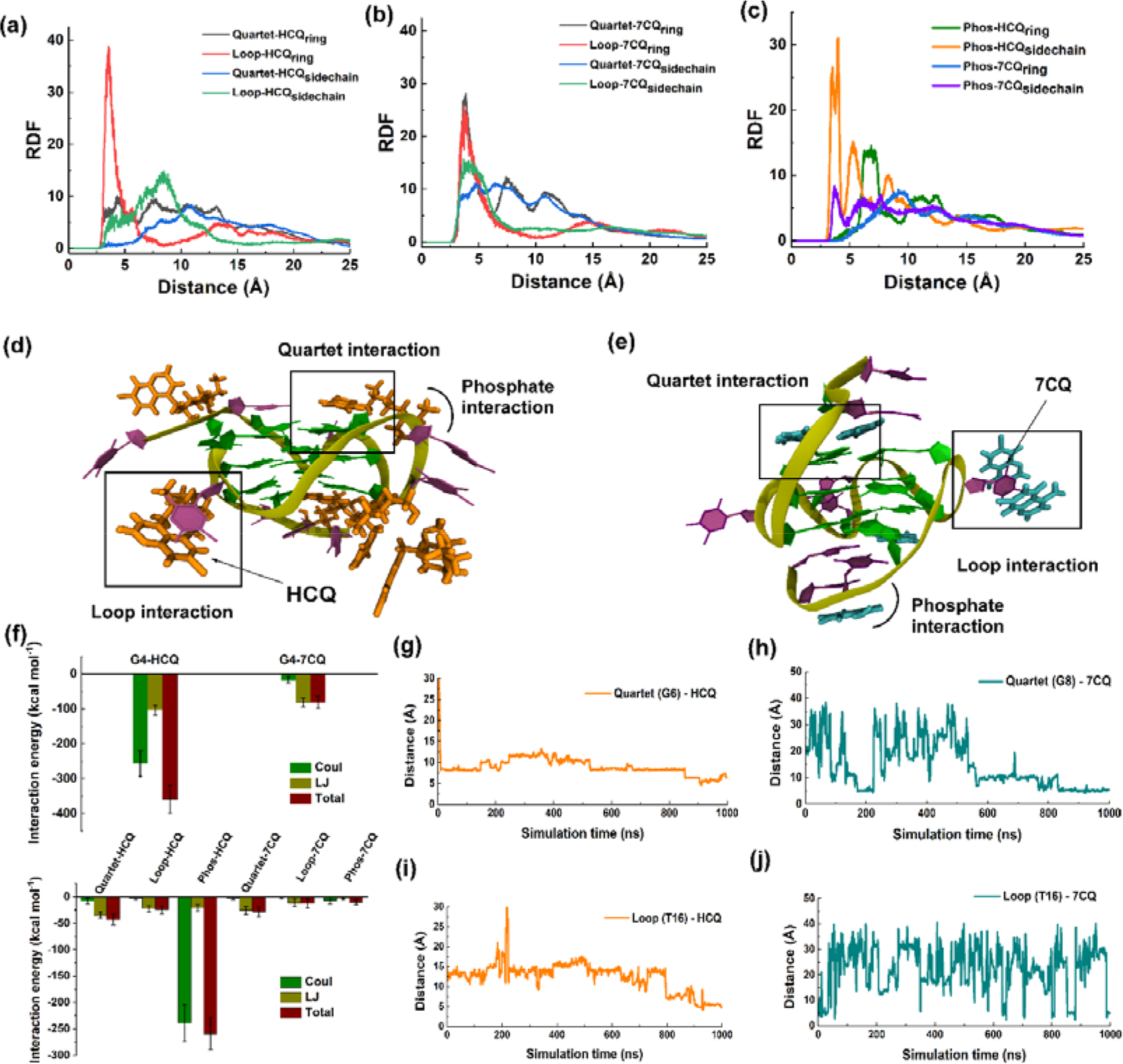
The radial distribution function (RDF) around the quartet and loop regions of G4 with (a) HCQ and (b) 7CQ ring as well as a side chain. (c) The RDF of the phosphate of the G4 with the ring as well as the side chain of the HCQ and 7CQ drugs. Snapshots of the G4 structure surrounded with (d) HCQ and (e) 7CQ molecules from the last frame of the simulation. The G bases in the G4 structure are shown in green color and the A, and T bases in the loop regions are shown in purple color. HCQ and 7CQ molecules are shown in orange color and cyan color, respectively. The stacking mode of interactions with G base in quartet and T base in loop regions were highlighted using a square box and the phosphate interaction is highlighted by an arrow. (f) The average of total interaction energy of the HCQ and 7CQ molecules with the whole c-myc G4 structure. The total interaction energy is dissected into coulombic (Coul) and Lennard-Jones (LJ) terms (upper panel). The total and dissected interaction energies of the HCQ and 7CQ with the quartet, loop, and phosphate backbone of G4 (lower panel). Error bars indicate the standard deviation (SD) of the data. The distance variation for the quartet and loop bases of G4 with HCQ (g, i) and 7CQ (h, j), respectively for the entire simulation time scale.

The interaction energy of the drug molecules with G4 depicts that the propensity of the interaction of HCQ with G4 is significantly higher than the 7CQ (Figure 4f, upper panel), in line with the binding data. It is also apparent that the positively charged HCQ interacts with G4 predominately through electrostatic interaction whereas the contribution of LJ is higher for neutral 7CQ. The interaction energy of the HCQ and 7CQ with the quartet, loop, and backbone of G4 suggest that the contribution of the LJ is higher for the interaction of the HCQ and 7CQ with the quartet and loop due to the stacking of the quinoline group of both the drug molecules on the quartet and loop regions of G4 (Figure 4f, lower panel). However, the contribution of the electrostatic interaction of HCQ with the phosphate backbone of G4 is significantly higher than 7CQ (Figure 4f, lower panel). The interaction energy of HCQ and 7CQ with different parts of G4 depicts that the electrostatic interaction between the HCQ and phosphate backbone is the main difference between the binding of the HCQ and 7CQ with G4. The time-dependent variation of the distance for the HCQ and 7CQ with the the guanine of the quartet and thymine of the loop of G4 indicate that binding of the HCQ is stable for the last 200 ns of the simulation time scale whereas 7CQ shows a higher degree of the fluctuation indicating that the less stable stacking of 7CQ with quartet and loop of G4 than HCQ (Figures 4 g-j). Finally, the interaction energy of the different quartets of G4 in the absence and presence of HCQ and 7CQ was estimated to understand the effect of the binding of these drugs on the stability of G4 (Figure S6). The interaction energy of the quartets is marginally higher on the binding of the G4 with HCQ than the case of 7CQ and without drug indicating that the binding of HCQ provides enhanced stability to G4 than the 7CQ and without drug cases (Figure S6). The MD simulation in corroboration of the NMR data implies that the electrostatic interaction between the side chain of HCQ and the phosphate backbone of the G4 anchor the enhanced stacking of the quinoline group of HCQ with the quartet and loop of G4 causing the increased stabilization of G4 induced by HCQ than without drug and 7CQ cases.

### HCQ-mediated selective stabilization of G4 Structures and transcriptional repression in cancer cells

The biophysical experiments revealed pronounced binding and stabilization of c-myc G4 structures by HCQ. qRT-PCR analysis of breast cancer MDA-MB 231 cells treated with varying concentrations of HCQ demonstrated a substantial reduction in c-myc mRNA levels. Notably, this reduction in c-myc mRNA level was observed even at low HCQ doses, with the most pronounced inhibition occurring at a concentration of 100 µM. These findings suggest that HCQ treatment is associated with a dose-dependent suppression of c-myc gene expression in MDA-MB 231 breast cancer cells, indicating a potential regulatory role for HCQ in modulating c-myc mRNA levels. (Figure 5a). Assessment of other genes harboring G4 structures (KRAS, c-kit, bcl2, VEGF) revealed HCQ-induced transcriptional repression, particularly notable for c-myc compared to modest effects on other genes (Figure 5b). Western blot analysis confirmed concentration-dependent decreases in c-myc protein levels following HCQ treatment, with a lesser impact on bcl2, requiring higher concentrations for comparable repression (Figure 5 c-d). Immunofluorescence analysis also corroborated HCQ-mediated reductions in c-myc protein expression in MDA-MB-231 cells (Figure 5e). Structure-activity relationship studies involving HCQ, CQ, Qn, and 7CQ exhibited marked suppression of c-myc expression by HCQ and CQ, while Qn and 7CQ had negligible effects, aligning with biophysical experiments and emphasizing the importance of side chain and cationic charge in selective interaction and stabilization of c-myc G4 (Figures 5 f-g, S7).

**Figure 5:**
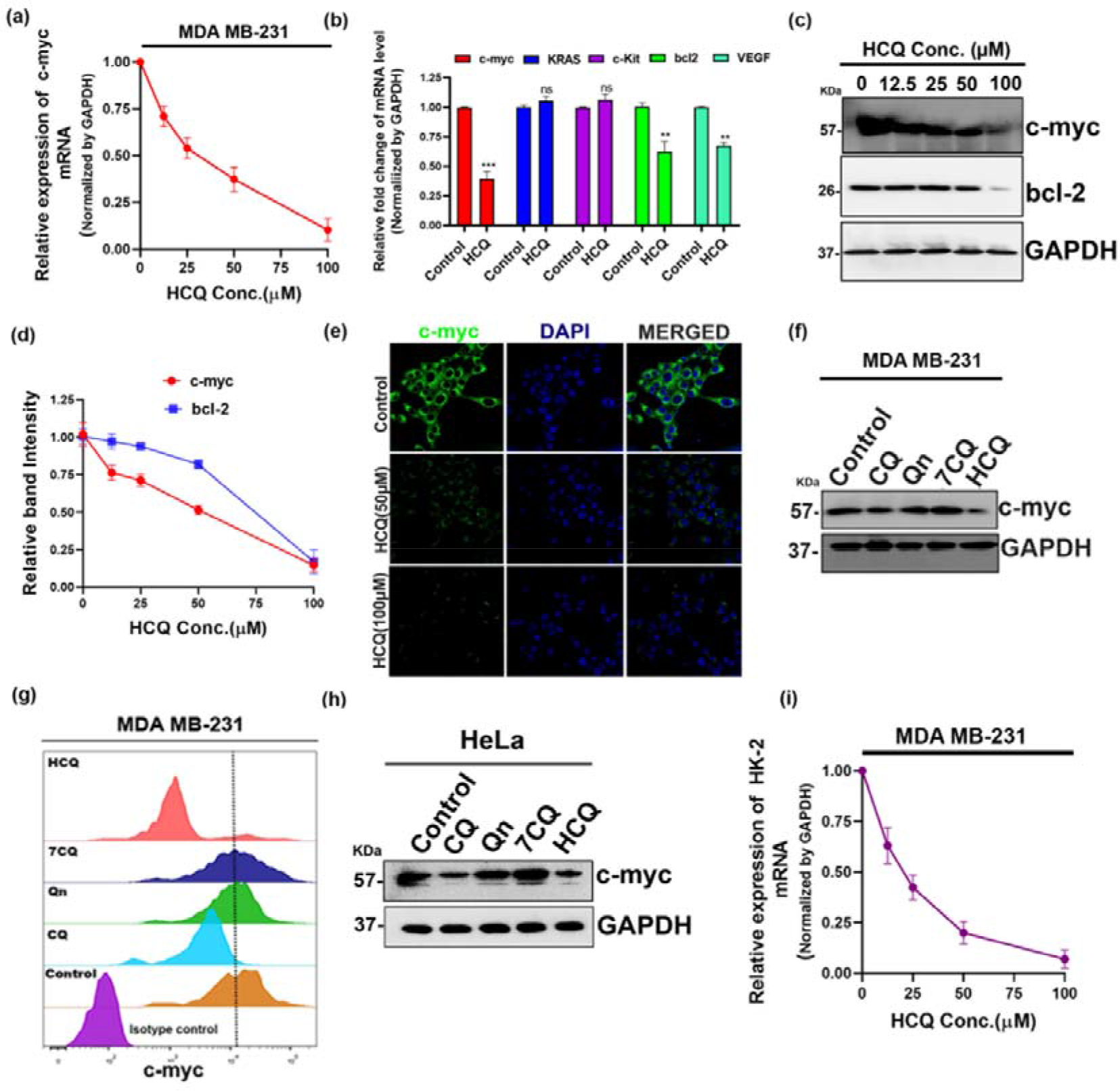
(a) qRT-PCR analysis of c-myc mRNA level following different doses of HCQ treatment for 48 hours in MDA-MB 231 cells. GAPDH was used as the normalization control. (b) qRT-PCR analysis of c-myc, KRAS, c-kit, bcl-2, and VEGF mRNA levels following 50 µM HCQ treatment for 48 hours in MDA-MB 231 cells. GAPDH was used as the normalization control. (c-d) Western blot analysis of c-myc and bcl-2 in MDA-MB-231 cells following HCQ treatments with different doses for 48 hours. The band intensity was calculated using ImageJ and plotted as relative band intensity using GraphPad Prism8. (e) Assessment of expression levels of c-myc via immunofluorescence in MDA-MB 231 cells following a 48-hour exposure to either 100 µM or 50 µM HCQ. Nuclei are counterstained in blue with DAPI. (f) Western blot analysis of c-myc expression in MDA-MB-231 cell line after the indicated treatments. (g) Flow cytometry analysis of c-myc expression in MDA-MB- 231 cells after the indicated treatments for 48 hours. (h) Western blot analysis of c-myc expression in HeLa cells following indicated treatments for 48 hours. (i) qRT-PCR analysis of HK-2 mRNA levels following dose-dependent HCQ treatment for 48 hours in MDA-MB 231 cells. GAPDH was used as the normalization control. The error bars indicate the mean ±SD of three independent experiments and the statistical significance levels denoted as follows: ns = not significant, P>0.05, *P<0.05, ** P<0.01, ***P<0.001, ****P<0.0001, assessed using ANOVA with Tukey post hoc test for multiple comparisons.

Further analysis in cervical cancer cells (HeLa) supported HCQ-induced c-myc suppression, suggesting a broad transcriptional inhibitory mechanism across cancer cell types (Figure 5 h, Figure S7). Although HCQ exhibited a slight decrease in bcl2 and VEGF levels, the extent was significantly less than that observed for c-myc. This decrease in bcl2 and VEGF transcriptional activity may be attributed to G4 stabilization or the indirect impact of reduced c-myc expression because c-myc is the designated transcription regulator for these two genes.^60^ *In-vitro* G4 stabilization studies indicated that HCQ’s effect on bcl2 is less pronounced than on KRAS and c-kit, whereas HCQ did not affect much to the transcriptional activity of KRAS and c-kit. Additional analysis of HK2, regulated by c-myc^61^ without G4 structures, also showed a significant decrease in HK2 mRNA levels post-HCQ treatment (Figure 5i). This raises the possibility that the reduction in bcl2 and VEGF transcriptional activity could be predominantly attributed to diminished c-myc expression directly influencing their transcription by binding to their promoters rather than G4 stabilization by HCQ. These findings emphasize HCQ’s selective and potent modulation of c-myc transcription with potential therapeutic implications.

### Anti-cancer activity of HCQ alone and in combination with doxorubicin in cell system and breast cancer murine model

Previous research has underscored the significant role of c-myc in governing cancer cell migration and invasion.^62^ In line with these insights, our investigation aimed to decipher whether HCQ treatment in breast cancer MDA-MB 231 cells influences these critical cancer cell characteristics. Employing transwell invasion and wound healing assays, a substantial reduction in both invasion and migration of triple-negative breast cancer (TNBC) cells following HCQ treatment was observed. Concurrently, viability assessments demonstrated a decrease in breast cancer cell viability upon HCQ treatment compared to vehicle-treated cells (Figure 5 a-c). Next, we explored the efficacy of HCQ, alone and in combination with doxorubicin (DOX), against breast cancer cells. Intriguingly, the combined treatment exhibited a more pronounced reduction in cell migration and invasion than either agent alone. MTT assay further unveiled a significant decrease in breast cancer cell viability with the DOX and HCQ combination, surpassing the effects observed with individual treatment. These findings hint at a potential synergistic interaction between HCQ and DOX in influencing breast cancer cell motility and survival, warranting further exploration in preclinical and clinical contexts. In summary, our study positions HCQ as a promising stand-alone agent for TNBC therapy through c-myc transcriptional downregulation and suggests its potential as an adjunctive in combination with chemotherapy (Figure 6 a-c).

**Figure 6:**
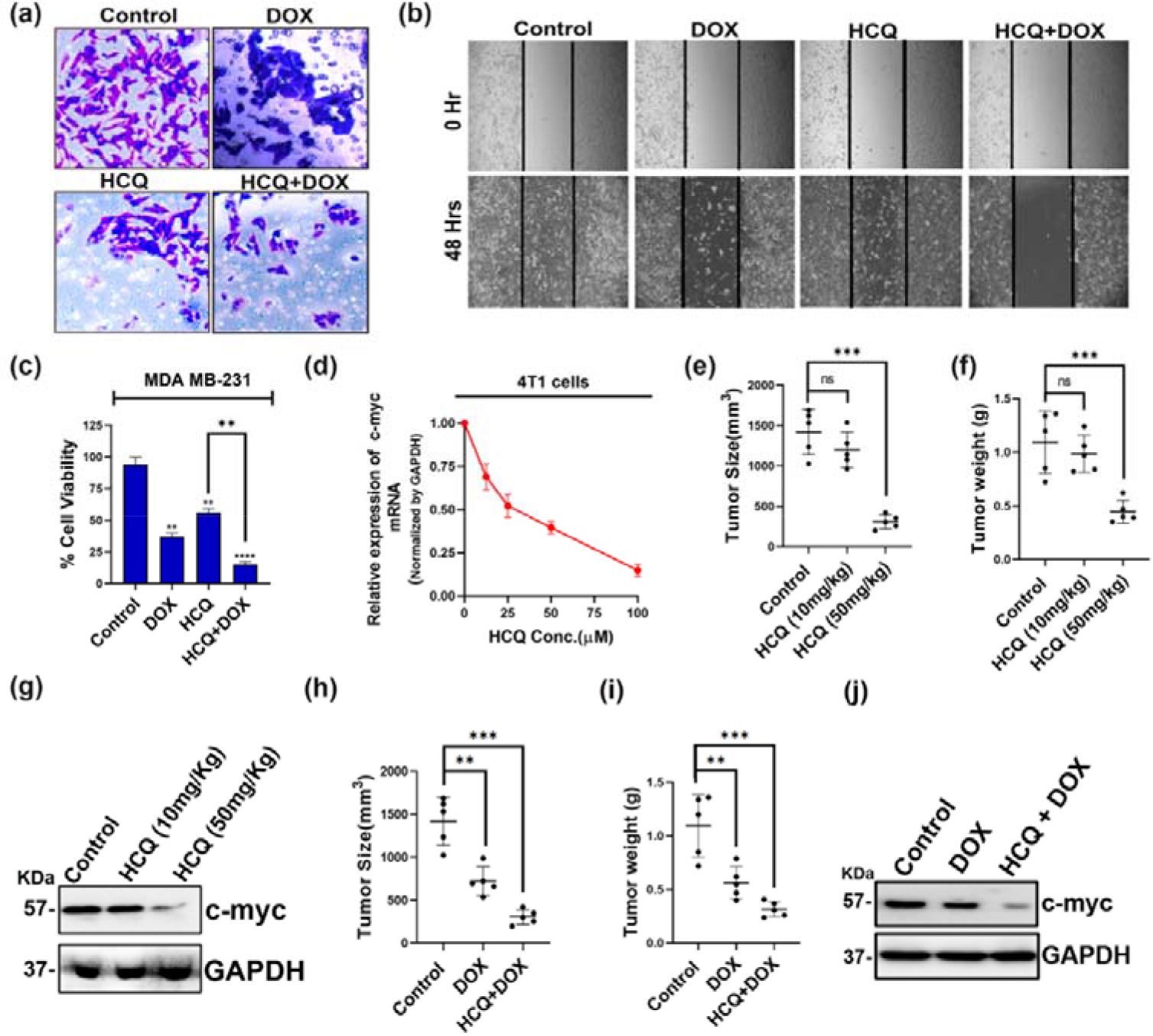
(a) Transwell invasion assay showing the extent of invasion of MDA-MB-231 cells after the indicated treatments (b) Representative images of wound healing assay showing the extent of migration of MDA-MB-231 cells after the indicated treatments. (c) Impact of HCQ, DOX, and combination therapy for 72 hrs. on the cell viability of MDA-MB-231 (Human triple-negative breast cancer) cells. Each experiment was conducted in triplicates, and the data were analyzed and plotted using GraphPad Prism 8. (d) qRT-PCR analysis of c-myc mRNA level following different doses of HCQ treatment for 48 hours in 4T1 cells. GAPDH was used as the normalization control. (e-f) statistical analysis of tumor size (e) and tumor weight (f) in Balb/c mice bearing tumor derived from 4T1 cells subjected to either control (saline) or HCQ (10mg/kgbdwt or 50mg/kgbdwt). (g) Western blot analysis of c-myc expression in tumors from BALB/c mice following the sacrifice. GAPDH was used as a loading control. (h-i) Measurement of tumor size (H) and tumor weight (I) in Balb/c mice following exposure to control, DOX, and combination therapy of both HCQ and DOX from 4T1-derived tumors. (j) Western blot analysis of c-myc expression in tumors following indicated treatments. The error bars indicate the mean ±SD of three independent experiments and the statistical significance levels denoted as follows: ns = not significant, P>0.05,*P<0.05, ** P<0.01, ***P<0.001, ****P<0.0001, assessed using ANOVA with Tukey post hoc test for multiple comparisons.

To unravel the *in-vivo* antitumor mechanism of HCQ, we conducted a preliminary investigation utilizing mouse-specific breast cancer (TNBC) cells (4T1). The study focused on assessing HCQ’s impact on c-myc transcription levels, revealing a concentration-dependent downregulation akin to its effects on human TNBC cells (Figure 6d). Subsequently, 4T1 cells were subcutaneously injected into Balb/c mice, and after tumor development, mice were treated with either a higher dose (50mg/kgbdwt) or a lower dose (10mg/kgbdwt) of HCQ. Results demonstrated a significant reduction in tumor volume and weight at the higher dose, while the lower dose proved ineffective. Western blot analysis indicated a substantial decrease in c-myc protein levels at 50mg/kgbdwt, suggesting a potential dose threshold for antitumor efficacy (Figures 6 e-g, S8). Interestingly, the lower HCQ dose, when combined with DOX, exhibited a significant reduction in tumor parameters compared to DOX alone, emphasizing potential combination therapy benefits. Furthermore, we analyzed the c-myc protein levels in tumor cells isolated from the treated mice. A significant reduction in c-myc levels was observed following Dox treatment and the reduction in the c-myc level is even more pronounced when HCQ and DOX are administered together (Figure 6 j, S8). These results suggest that the HCQ downregulates the c-myc transcription possibly by binding with the G4 sequence of the c-myc gene, which reduces tumor size and volume. This study indicates the potential of HCQ in the treatment of breast cancer either alone or adjunctive therapeutic manner.

## Discussion

In the present study, the molecular picture of the binding of the FDA-approved drugs HCQ and CQ with c-myc G4 and its effect on gene regulation and tumor progression using the *in-vitro* and *in-vivo* biological methods have been demonstrated. The fluorescence binding data indicate that HCQ and CQ both have the capability of binding with G4 structures associated with different oncogenes. The comparison of the binding data of HCQ, CQ, and its analogs Qn and 7CQ indicate that the charged group and the side chain of the HCQ and CQ enhance the binding tendency of these drugs with G4 structure compared to their analogs Qn and 7CQ. The melting temperature data of G4 with HCQ and CQ indicate that the binding of HCQ and CQ stabilizes the G4 structure of the c-myc oncogene significantly higher than others. The molecular level information of the binding of the HCQ with the G4 structure has been speculated using the ^1^H-NMR in the imino region of the G4 and MD simulation. The NMR and MD simulation data suggest that electrostatic interaction between the side chain of HCQ and the phosphate backbone of the G4 anchors the enhanced stacking of the quinoline group of HCQ with the quartet and loop of G4. There are earlier reports in which the ligand has been found to bind with the G4 by stacking at one end of the G4.^57, 59^ The melting temperature CD data indicate that the binding of HCQ and CQ profoundly stabilizes the G4 associated with the c-myc oncogene. HCQ-mediated downregulation of c-myc also correlates with reduced invasion and metastasis of breast cancer cells signifying its anticancer properties as a stand-alone medicine. We also investigated whether HCQ can enhance the efficacy of DOX, a standard treatment regimen for breast cancer. The *in-vitro* experiments showed a significant reduction of invasion and metastasis of breast cancer cells when the cells were treated with both HCQ and DOX with respect to the cells treated with DOX alone. Therefore, the study revealed that the combination of HCQ with the frontline chemotherapeutic drug such as DOX augments its antineoplastic activity beyond either single agent treatment. Furthermore, a pre-clinical allograft murine model was employed to validate cell-based observation in an *in-vivo* setting. The data indicate that higher doses of HCQ as a monotherapy sufficiently reduce tumor growth; however, comparatively lower HCQ doses are only effective in restricting tumor progression when combined with DOX treatment.

There are few studies earlier in which the probable action of the HCQ and CQ on the cancer cells has been discussed. It has been proposed that HCQ and CQ may suppress immune function and inhibit autophagy causing the death of the tumour cells.^38, 63^ However, in another study, it has been proposed that HCQ decreases the benefit of anti-PD1-immune checkpoint blockade in tumor immunotherapy.^39, 43, 44^ CQ has been proposed to affect the TLR9-mediated NF-kB signaling pathway based on the in-vitro experiments, however, it could not show the desired effect during the in-vivo experiments.^41^ CQ has also been proposed to regulate p53 protein and activate the p53-dependent transcription of pro-apoptotic genes. In contrast, few reports indicate that CQ induces an anticancer effect independent of the p53 pathway and status.^42, 64^ In contrast to all these studies, we have illustrated that these drugs recognize the c-myc G4 and downregulate the mRNA and protein levels associated with the gene as well as decrease the tumor size, intrinsically as well as in combination modes with chemotherapeutic drugs.

## Conclusion

A variety of synthetic small molecules have been reported to bind to the c-myc G4, however, these molecules could not be used as drug molecules due to the poor biocompatibility and pharmacological issues. In this regard, our work demonstrates the capability of the known FDA-approved immunomodulator drugs HCQ and CQ which are in the preclinical trial stage for the cancer to target specifically the c-myc expression through direct G4 DNA interaction. The recognition of c-myc-G4 with the drug is mainly triggered by the electrostatic interaction between the charged side chain of the HCQ with the phosphate backbone of G4 which anchors the stacking of the quinoline ring of HCQ on the quatret and loop of G4. Beyond intrinsic anti-cancer effects, HCQ may also re-sensitize cells to chemotherapy that would ordinarily upregulate pro-survival c-myc levels. Our data advocate further preclinical development of this approved drug for potential clinical benefit in breast cancer care.

## Supporting Information

Spectroscopy data of the binding, and melting of different G4 sequences with drug molecules, NMR data of the binding of c-myc G4 with 7CQ, The MD simulation data of the G4 structure of c-myc with HCQ and 7CQ, quantification of the *in-vitro and in-vivo* biological experiments.

## Acknowledgments

PCS thanks DST-SERB (CRG/2019/001389) for financial support. AC and SP thank IACS for their fellowship. AB thanks DST-Inspire for the fellowship.

## Table of Contents

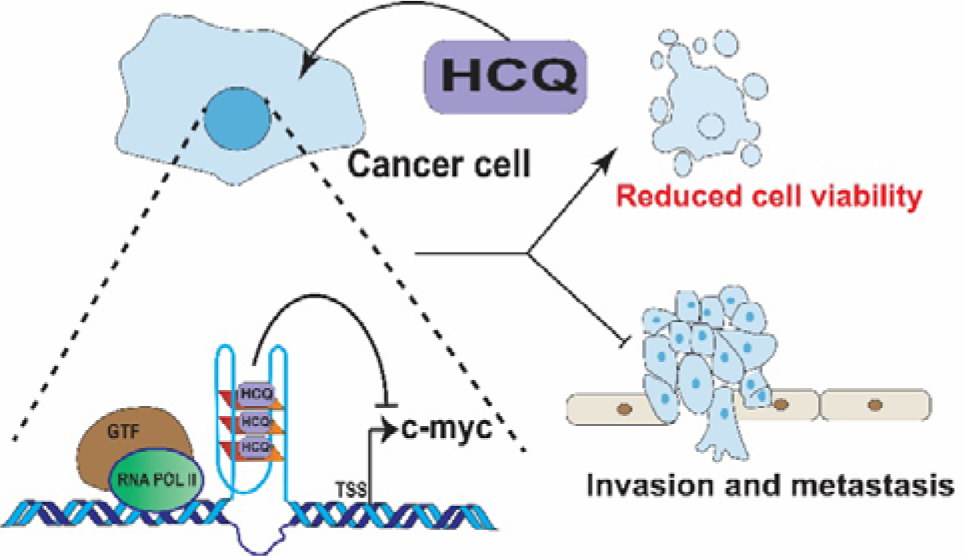

