## supporting for "G-Quadruplex Mediated c-myc Specific Downregulation: A Unique Pathway of the Anticancer Action of Immunomodulator Drugs"

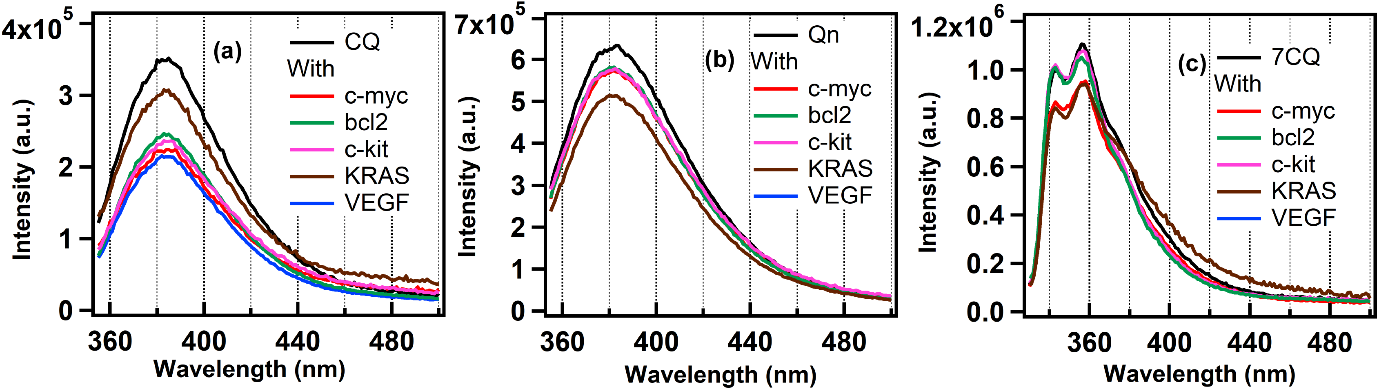

**Figure S1:**The emission spectra of CQ (2µM, a), Qn (2µM, b), 7CQ (2µM, c) in buffer and different G4 DNA (2µM) sequences.

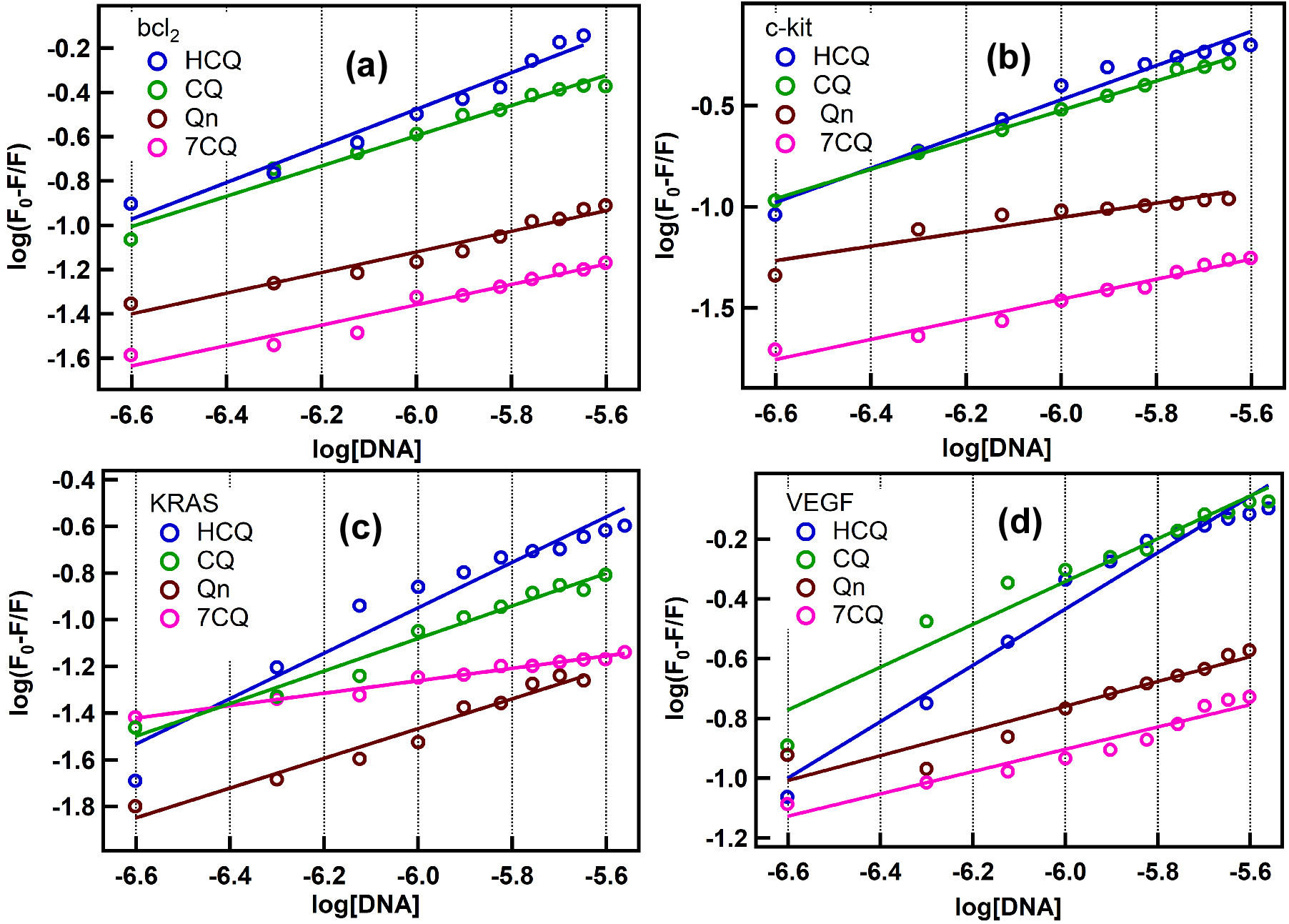

**Figure S2:** The plot of the change in the intensity of the emission maxima of HCQ, CQ, and its analogs Qn and 7CQ with respect to the concentration of bcl2 (a), c-kit (b), KRAS (c), and VEFG (d) DNA sequences, respectively. The data has been fitted with the modified Stern-Volmer equation to calculate the binding constant value of the drug with G4 sequences.

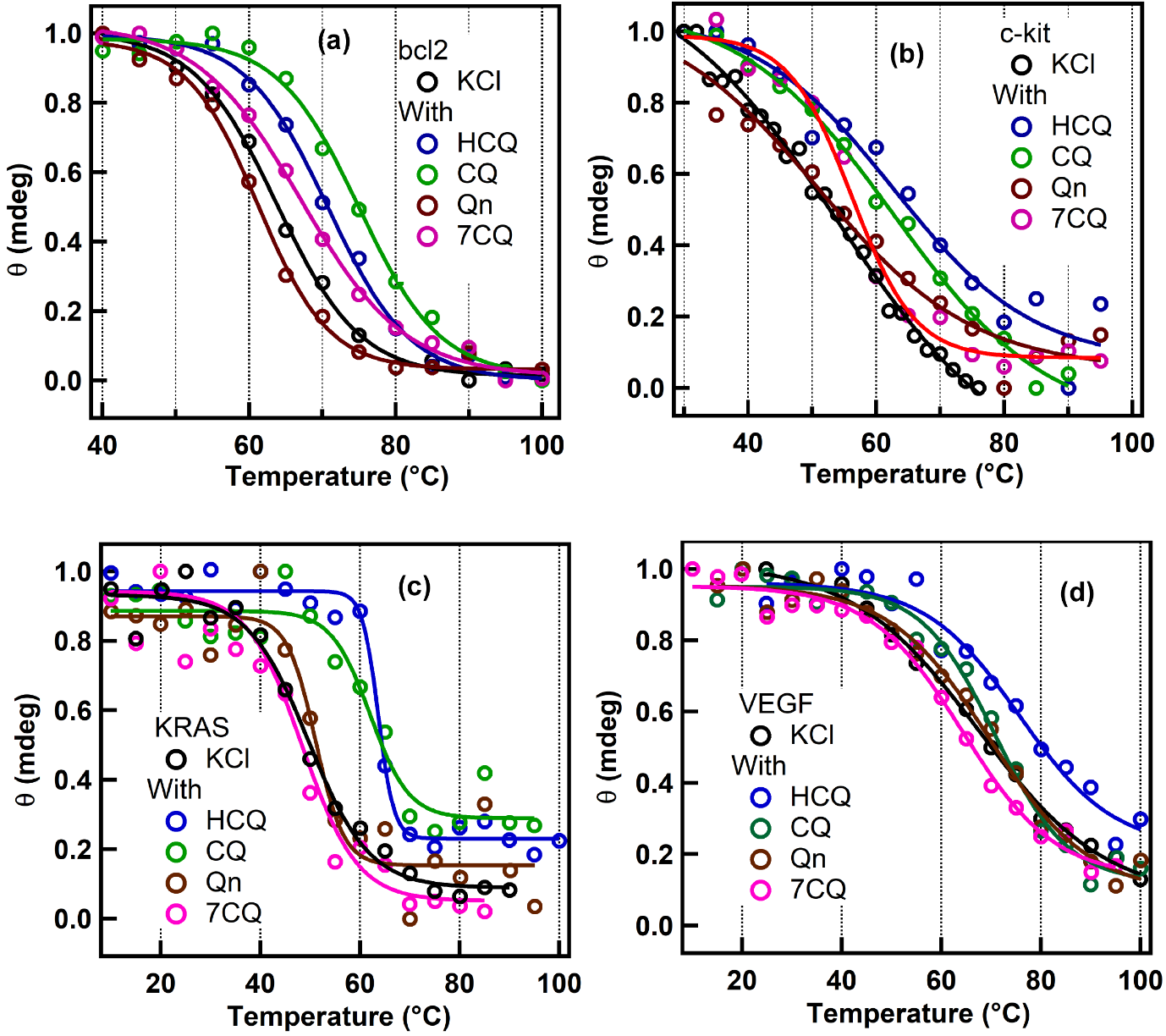

**Figure S3:**The melting curve of the bcl2 (5µM, a), c-kit (5 µM, b), KRAS (5 µM, c), VEGF (5 µM, d) in KCl and the presence of different drug molecules (50 µM each).

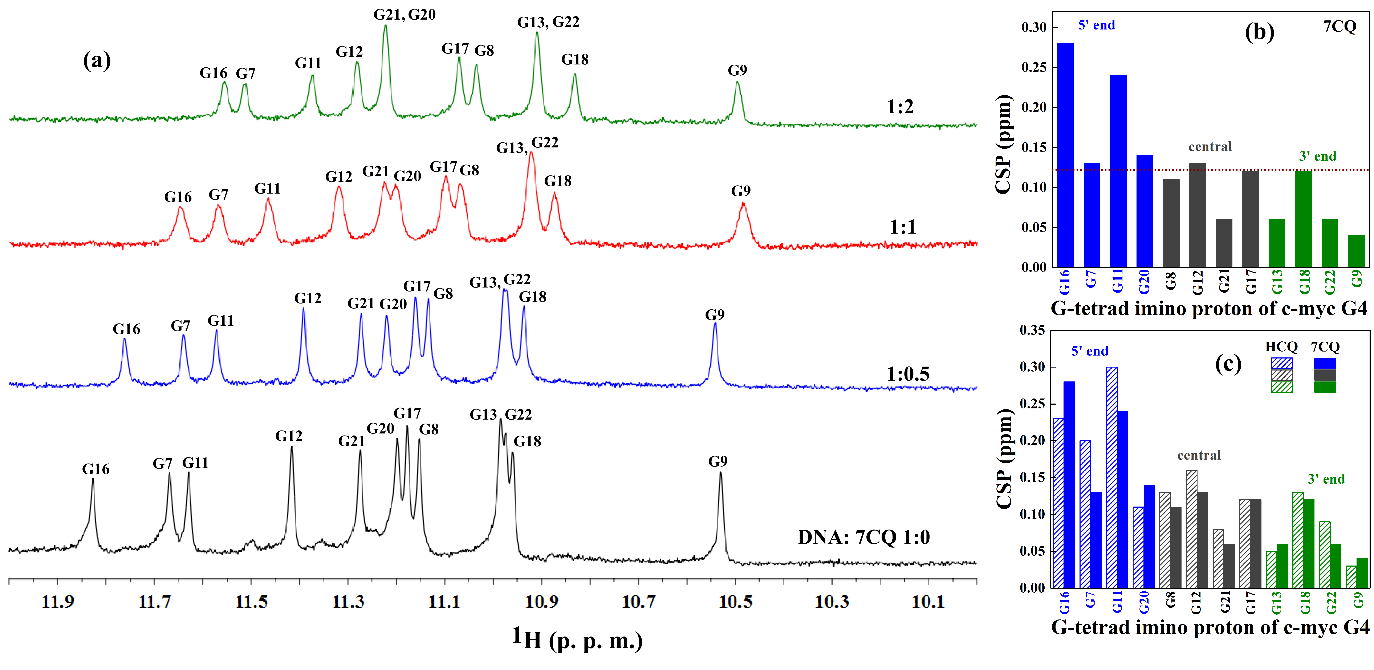

**Figure S4:** (a)^1^H-NMR spectra representing the imino region of the G4 during the titration of c-myc with 7CQ. The G4-imino protons of tetrad and the ratio of c-myc and HCQ are labelled in each spectrum. (b) The plot of chemical shift perturbation for (CSP) each imino protons of c-myc G4 tetrads at 5’ (blue end), central (black), and 3’ (green) terminals in the presence of 7CQ. (c) The comparison of the change of CSP values of c-myc G4 in the case of HCQ and 7CQ, respectively.

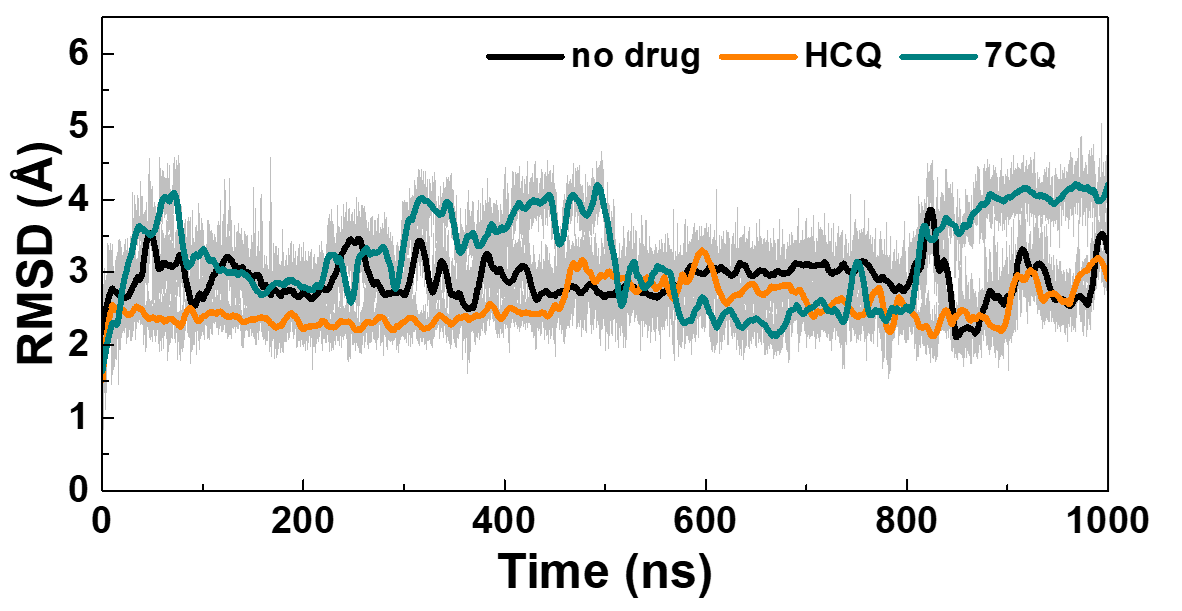

**Figure S5.** The root mean square deviation (RMSD) of G4 structure in the absence of drug (no drug) and in the presence of HCQ and 7CQ for the whole simulation time.

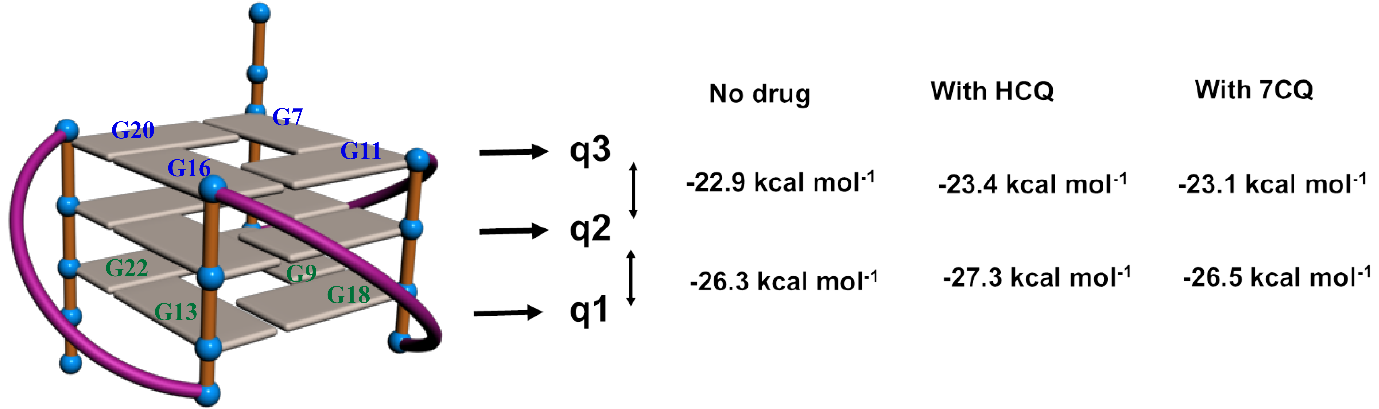

**Figure S6.** Schematic of the parallel G4 structure of c-myc with the representation of G bases involved in quartet 1 (q1), quartet 2 (q2), and quartet 3 (q3). The interaction energy between q1-q2 and q2-q3 in the absence of the drug and in the presence of HCQ and 7CQ were represented. The atoms of the rings of the G bases involved in the quartet as shown in the schematic were selected for the interaction energy calculation.

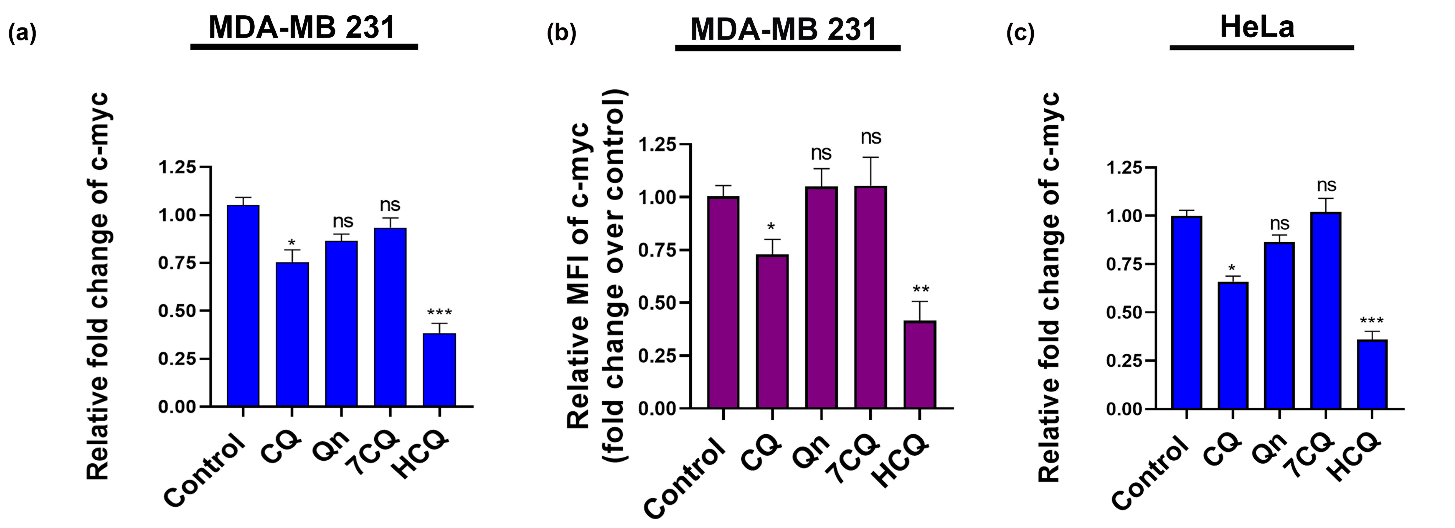
**Figure S7:** (a) Densitometric analysis of Western blot of c-Myc expression in MDA-MB-231 cell line after treatment with CQ, Qn, 7CQ and HCQ (dissolved in pH 7 buffer) for 48hrs using ImageJ. (b) Quantification of the relative MFIs obtained from the flow cytometry analysis of c-Myc expression after drug treatments. (c) Densitometric analysis of Western blot of c-Myc expression in HeLa cell line after treatment with CQ, Qn, 7CQ and HCQ(dissolved in pH 7 buffer) for 48 hrs using ImageJ. All the graphs were generated using GraphPad Prism8, presenting the combined (mean) outcomes derived from three independent experiments. The error bars indicate the range of variability observed across these repetitions (mean ± standard deviation, n=3), with statistical significance levels denoted as follows: ns = not significant, P>0.05, *P<0.05, ** P<0.01, ***P<0.001, ****P<0.0001, assessed using ANOVA with Tukey post hoc test.

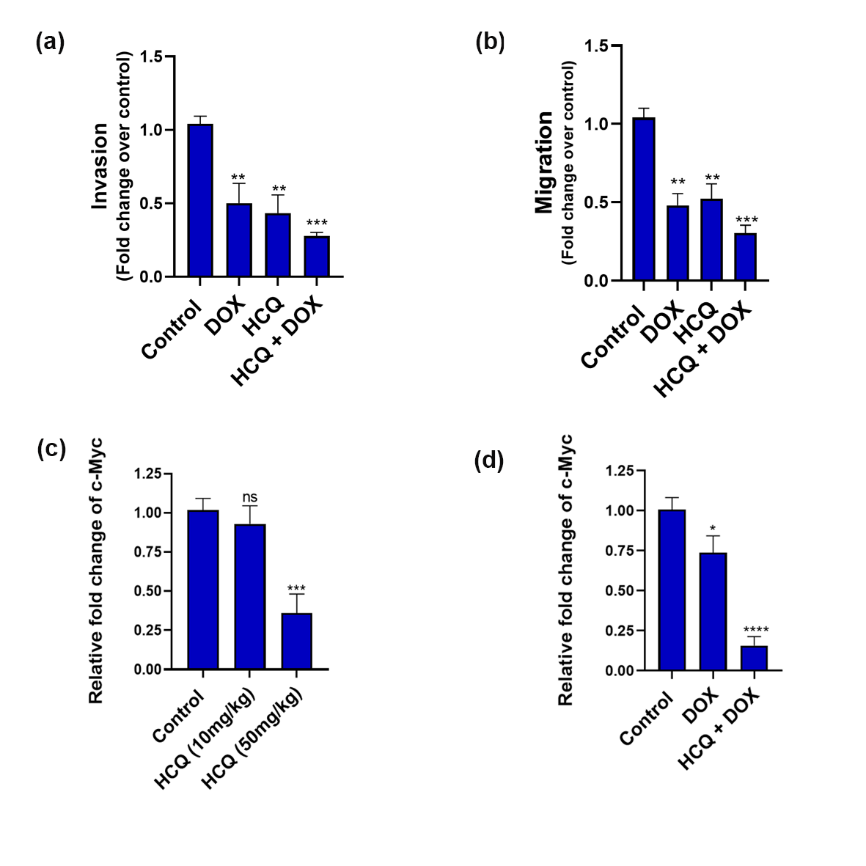

**Figure S8:** (a) Quantification of Transwell invasion assay in MDA-MB-231 cells after the indicated treatments. (b) Quantification of wound healing assay showing the extent of migration of MDA-MB-231 cells after the indicated treatments. (c)Densitometric analysis of Western blot of c-myc expression in tumors from BALB/c mice following the sacrifice. GAPDH was used as a loading control. (d) Densitometric analysis ofWestern blot data of c-myc expression in tumors following indicated treatments. The error bars indicate the mean ±SD of three independent experiments and thestatistical significance levels denoted as follows: ns = not significant, P>0.05, *P<0.05, ** P<0.01, ***P<0.001, ****P<0.0001, assessed using ANOVA with Tukey post hoc test for multiple comparisons.

**Table S1: Sequence of primers for qRT-PCR analysis**

| **Gene name** | **Forward(5`-3`)** | | **Reverse (5`-3`)** |
| --- | --- | --- | --- |
| **c-myc** | CCTGGTGCTCCATGAGGAGAC | CAGACTCTGACCTTTTGCCAGG | |
| **bcl2** | ATCGCCCTGTGGATGACTGAGT | GCCAGGAGAAATCAAACAGAGGC | |
| **KRAS** | CAGTAGACACAAAACAGGCTCG | TGTCGGATCTCCCTCACCAATG | |
| **c-kit** | CACCGAAGGAGGCACTTACACA | TGCCATTCACGAGCCTGTCGTA | |
| **VEGF** | TTGCCTTGCTGCTCTACCTCCA | GATGGCAGTAGCTGCGCTGATA | |
| **HK2** | GAGTTTGACCTGGATGTGGTTGC | CCTCCATGTAGCAGGCATTGCT | |
